# Promoter-specific changes in initiation, elongation and homeostasis of histone H3 acetylation during CBP/p300 Inhibition

**DOI:** 10.1101/2020.09.26.315002

**Authors:** E Hsu, NR Zemke, AJ Berk

## Abstract

Regulation of RNA Polymerase II (Pol2) elongation in the promoter proximal region is an important and ubiquitous control point for gene expression in metazoan cells. We report that transcription of the adenovirus 5 E4 region is regulated during the release of paused Pol2 into productive elongation by recruitment of the super elongation complex (SEC), dependent on promoter H3K18/27 acetylation by CBP/p300. We also establish that this is a general transcriptional regulatory mechanism for ∼6% of genes expressed with FPKM>1 in primary human airway epithelial cells. We observed that a homeostatic mechanism maintains promoter, but not enhancer H3K18/27ac in response to extensive inhibition of CBP/p300 acetyl transferase activity by the highly specific small molecule inhibitor A-485. Further, our results suggest a function for BRD4 association at enhancers in regulating paused Pol2 release at nearby promoters. Taken together, our results uncover processes regulating transcriptional elongation by promoter region histone H3 acetylation and homeostatic maintenance of promoter, but not enhancer, H3K18/27ac in response to inhibition of CBP/p300 acetyl transferase activity.

## Introduction

In addition to RNA polymerase II (Pol2) pre-initiation complex (PIC) assembly and initiation, the transition from promoter-proximal paused Pol2 to productively elongating Pol2 is an essential step in gene transcription and an important process in the overall multi-component orchestration of gene expression (1–3). After the recruitment of Pol2 to a promoter by its general transcription factors and assembly of a PIC (4, 5), transcription initiation occurs concurrently with TFIIH phosphorylation of Ser5 of the Pol2 heptapeptide repeat C-terminal domain (CTD) (6). In metazoan cells, Pol2 then transcribes approximately 30-60 bases downstream of the transcription start site (TSS) and pauses because it is bound by negative elongation factor (NELF) and DRB-sensitivity inducing factor (DSIF, Spt4 and Spt5 in *S. cerevisiae*) (7, 8). Recruitment of P-TEFb and its enzymatic subunits CDK9-Cyclin T results in the phosphorylation of NELF, DSIF, and Ser2 of the Pol2 CTD, whereupon NELF dissociates and Pol2 is released and proceeds to productive elongation (6–9).

Histone acetylation is well known to contribute to a permissive chromatin state for Pol2 PIC assembly at active promoters, and there is recently published work concerning its function in facilitating transcriptional elongation as well. For example, the chromatin reader protein BRD4 is thought to recruit P-TEFb (CDK9-Cyclin T) to promoters and serves as a Pol2 elongation factor dependent on its interactions with acetylated histone lysines through its bromodomains (10). In addition, H3 acetylation mediated by the Drosophila CBP ortholog stimulates productive elongation past the +1 nucleosome (11). Recruitment of the yeast histone chaperone FACT by acetylated H3 has also been shown to stimulate elongation (12).

The SEC is a multi-subunit complex comprised of P-TEFb (CDK9-Cyclin T) along with AF4/FMR2 proteins AFF1/4, ELL family members ELL1/2/3, ELL-associated factors EAF1/2, and one or the other highly homologous proteins AF9 or ENL containing YEATS acetyl-lysine-binding domains (13). There are various forms of the SEC, including SEC-like complexes that contain different combinations of elongation factors suggesting diversity in their regulatory mechanisms (13). P-TEFb, a central serine/threonine-kinase, an AFF scaffold protein, and ENL or AF9 are consistent components of SEC complexes. ENL and AF9 have been functionally linked to SEC recruitment to acetylated chromatin via their YEATS domains (14, 15). The SEC then stimulates transcription elongation through interactions with the PAF1 complex (16), which blocks NELF-binding to Pol2 (6), and DOT1L, which deposits the active chromatin modification H3K79me in the first intron (14, 17). Importantly, AF9 and ENL YEATS domains bind to active chromatin marks H3K9ac and H4K15ac (15), and to a lesser extent, H3K18/27ac (14), and are essential for SEC-dependent activation of a luciferase reporter driven by the HIV-1 LTR (16). Despite these conclusions, the function of histone acetylation during the transition from promoter-proximal paused to productively elongating Pol2 remains incompletely understood.

We previously reported that p300/CBP acetylation of H3K18 and K27 in the two to three nucleosomes spanning the transcription start site (TSS) had very different effects on distinct steps in transcription from different human adenovirus 5 (HAdV-5) early promoters (19). At the E3 promoter, loss of H3K18/27ac in the promoter region had little effect on PIC assembly, and the rate of E3 mRNA synthesis was only modestly reduced (<2-fold) compared to transcription activated by wt E1A which induces H3K18/27ac at the early viral promoters. In contrast, PIC assembly at the E2early promoter was almost eliminated by loss of promoter H3K18/27ac, and E2 mRNA synthesis was undetectable at 12 h p.i. (19). For E4, loss of promoter H3K18/27ac had little effect on PIC assembly, but caused a significant (∼10-fold) decrease in E4 transcription at 12 h p.i. (19). This result was particularly striking as it suggested that E4 transcription is regulated by promoter H3K18/27 acetylation at a step in transcription subsequent to PIC assembly, possibly during release of promoter-proximal paused Pol2.

To investigate the function of H3K18/27ac in transcriptional elongation at E4, we mapped the association of transcriptionally active Pol2 on the Ad5 genome using GRO-seq (Global Run-On sequencing) (1). We found defective paused Pol2 release at E4 in cells expressing an E1A mutant (“E1A-DM”) with polyalanine substituted for two highly acidic regions of the E1A activation domain (AD) that each mediate an interaction with p300/CBP (19). ChIP-seq for BRD4 and SEC components CDK9, AF9, and ENL revealed decreased SEC recruitment to E4 by E1A-DM compared to wt E1A. Using the specific small molecule inhibitor of CBP/p300 acetyl-transferase activity A-485 (20), we determined that CBP/p300 HAT activities are essential for maximal paused Pol2 release and SEC recruitment at the E4 promoter, but not at the E3 promoter.

We then extended our studies to the human genome, where we found that 2 h of A-485 treatment resulted in hypoacetylation of total cell H3K18/27 to a new, hypoacetylated steady-state. This was associated with defective pause-release at a subset of active genes (∼6%) where promoter H3K18/27ac was decreased by the drug. Differences in the sensitivity of transcription from different promoters to H3K18/27ac correlated with differences in SEC component association with the genes after A-485 treatment. This was similar to what we had observed for the HAdV-5 E4 promoter during activation by the multi-site E1A mutant (DM-E1A) with mutations in the E1A-AD acidic peptides required for p300/CBP binding to the E1A-AD. We also found that at a subset of enhancers with greatly decreased H3K18/27ac in response to A-485 treatment, H3K9ac is sufficient for BRD4 binding and stimulation of Pol2 pause-release. Based on these results, we propose mechanisms of BRD4 and SEC recruitment by histone H3 acetylation during the transition from promoter-proximal paused to productively elongating Pol2, and report a homeostatic process that maintains promoter H3K18/27ac.

## Results

### CBP/p300 acetylation of promoter histone H3K18 and K27 stimulates paused Pol2 release at the human adenovirus 5 E4 promoter, but is not required at the E3 promoter

Transcription from human adenovirus 5 (HAdV-5) early promoters is activated by the first viral proteins expressed following infection, the E1A isoforms, primarily large E1A (Figure S1A). While transcriptional activation from the viral early promoters is entirely dependent on the interaction of the large E1A isoform with the mediator of transcription complex, transcription of E4 is stimulated an additional ten-fold through interactions between CBP/p300 and two highly acidic regions immediately flanking the E1A mediator-binding region (large E1A aa residues 133—138 and 189—200 Figure S1(a,b) (19). Separate Ad5 expression vectors were constructed that express the wt E1A region from the wt E1A promoter/enhancer region, or DM-E1A with several mutations that convert these acidic peptides in wt E1A to polyAla (Fig S1b).

To analyze the effects of promoter H3K18/27ac on Pol2 elongation through the early Ad5 genes, we applied the GRO-seq method, which reveals the position and direction of transcribing Pol2 by BrU-labeling of 3’-ends of nascent RNA transcripts in isolated nuclei (1). The nuclei were first washed with the non-ionic detergent Sarkosyl to remove proteins from chromatin that block transcription elongation and to prevent Pol2 initiation, so only actively transcribing RNA polymerases at the time the nuclei were isolated produce BrU-labeled RNA (1). To avoid possible effects of cellular mutations in stable cell lines, we performed these studies in primary human bronchial-tracheal epithelial cells (HBTECs) derived from human adult lung transplant donors. These HBTECs are a cell culture model for the airway epithelial cells infected by HAdV-5 in humans.

We infected HBTECs with the wt E1A vector, and, separately, the DM-E1A vector expressing mutant E1A with polyAla substitutions of the two highly acidic peptides flanking CR3 (see Figure S1b). Wt E1A binds CBP/p300 through these highly acidic peptides, inducing histone H3 acetylation at K18 and K27 by the CBP/p300 acetyl transferase domain, in the viral E2early, E3, and E4 promoter regions (19). In cells expressing DM-E1A, which does not interact in vivo with p300 though the E1A activation domain (E1A-AD) (19), there was far less H3K18/27ac in these early viral promoter regions (Figure 1a). To express equal steady-state levels of the wt E1A and less stable DM-E1A proteins, infections were performed at a higher multiplicity of infection for the DM-E1A vector than for the wt E1A vector (19) (Figure S1(c)). Cells infected with the wt E1A vector were also co-infected with sufficient E1A deletion mutant *dl312* to maintain the same number of viral DNA molecule templates for the viral early regions (∼100 viral DNA molecules per nucleus) in cells expressing the same level of wt E1A and the DM-E1A protein (19) (Figure S1(c)).

**Figure 1:**
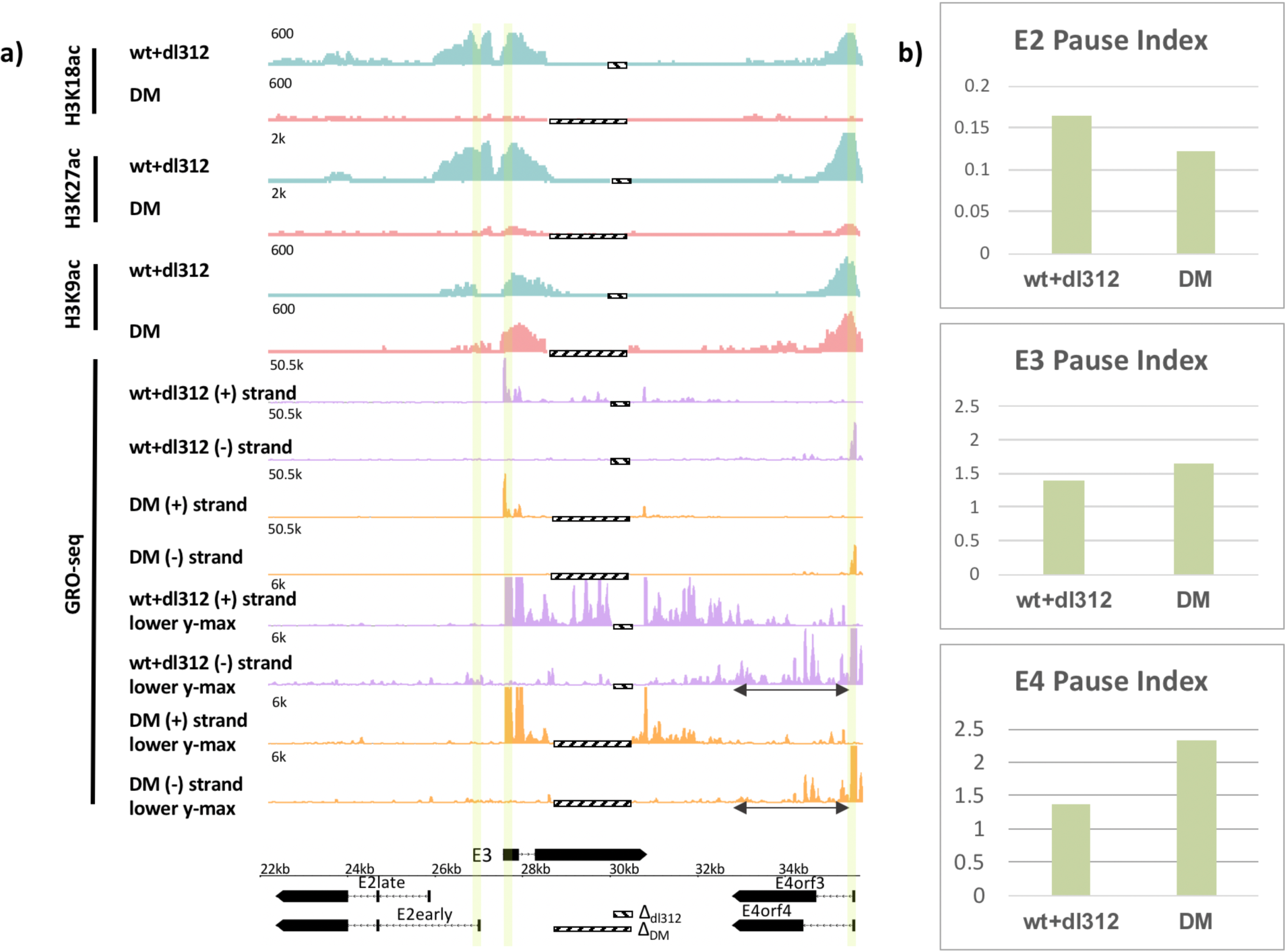
Promoter H3K18/27 acetylation activated by E1A-AD–CBP/p300 interactions stimulates paused Pol2 release at adenovirus promoter E4. **(a)** (bottom) map of the major HAdV-5 early E2, E3 and E4 mRNAs. Deletions in the E3 regions of *dl312* and the E1A-DM vector are shown by cross-hatched horizontal bars. Vertical stripes highlighted in yellow indicate promoter proximal regions. GRO-seq counts from primary HBTECs infected with wt+dl312 or DM vectors at 12 h post-infection (p.i.), were plotted on the Ad5 genome with H3K18ac, H3K27ac and H3K9ac ChIP-seq data (19). GRO-seq tracks are shown for the two viral DNA strands (+, transcribed to the right; and –, transcribed to the left), with two different y-axis scales to allow visualization of high and low amplitude peaks. The double-headed arrows in the GRO-seq plots in the E4 region refer to gene body regions discussed in the text. **(b)** Pause indexes for E2, E3, and E4 in cells expressing wt E1A or DM-E1A. Pause index is the ratio of: reads in the promoter region (TSS to +200) to reads in the gene body (+200 to TTS).

GRO-seq data at 12 hours post infection (h pi) with the vector expressing wt E1A revealed peaks of paused Pol2 with the expected orientation and location of promoter-proximal paused Pol2, ∼40-60 bp downstream from the E3 and E4 TSSs (Figures 1a, highlighted, and 2a). At 12h pi, very low GRO-seq signal was observed at the E2 early promoter or within the E2 gene body in wt E1A expressing cells compared to E3 and E4 (Figure 1a). This was probably because E2early transcription is delayed compared to E3 and E4 in these primary cells, and increases by 18 h p.i. (19). The low GRO-seq signal in the E2early promoter region and gene body was decreased further in cells expressing DM-E1A compared to wt E1A (Figure 1a), supporting our previous conclusion that E2early transcription is regulated by H3K18/27ac in the promoter region because it is required for rapid PIC assembly (19).

**Figure 2:**
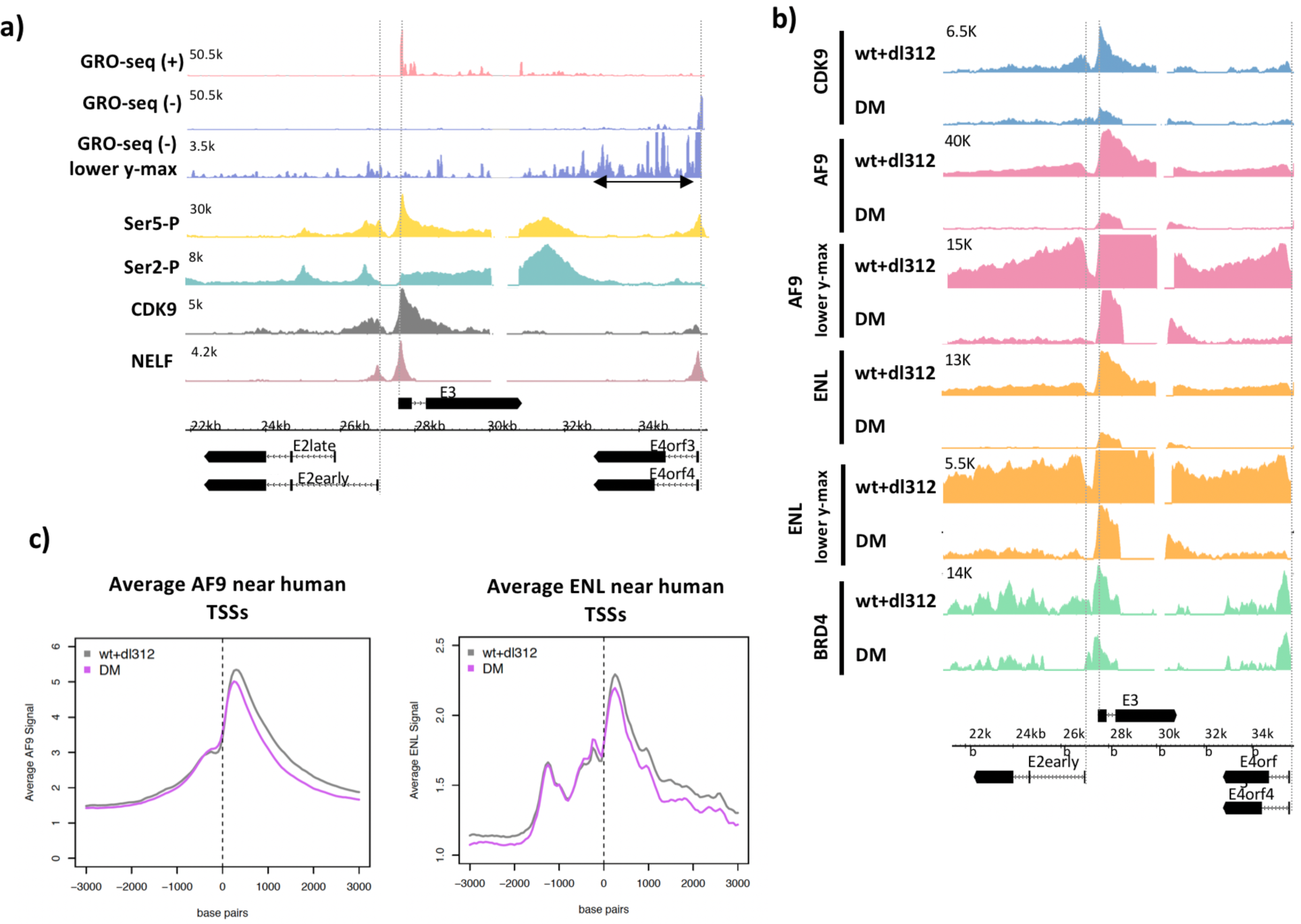
Ser5-P, Ser2-P, CDK9, NELF, and SEC subunits on the Ad5 genome. (a) Ser5-P, Ser2-P, CDK9, and NELF ChIP-seq plotted with GRO-seq in cells expressing wt E1A. (b) CDK9, AF9, ENL, and BRD4 ChIP-seq in cells expressing wt or DM E1A. AF9 and ENL ChIP-seqs are plotted with 2 different y-axes. (c) Average plots of AF9 and ENL ChIP-seq counts near TSSs on the human genome.

To determine the degree of promoter-proximal pausing in the E3 and E4 promoter regions where Pol2 association is detected by ChIP-seq at 12 h pi (19), we calculated the Pol2 pausing index (PI, (1)). The PI equals the number of GRO-seq reads in the promoter-proximal region (TSS to +200 bp) divided by the total GRO-seq reads in the gene body (+201 to TTS). The GRO-seq reads in the promoter-proximal region reflect the amount of promoter-proximal paused Pol2 at the time the nuclei were isolated, while the GRO-seq reads in the gene body reflect the amount of elongating Pol2 subsequent to pause-release. Therefore, an increase in a gene’s PI indicates a reduced rate of promoter-proximal pause-release.

After activation by DM-E1A, the PI at E4 increased almost 2-fold compared to E4 transcription activated by wt E1A (wt PI=1.37 vs. DM PI= 2.33) (Figure 1b). The vectors expressing wt E1A and DM-E1A had different size deletions in E3 due to the details of their constructions (Figure 1a, bottom), but the calculation of PI for E3 was based on the regions of E3 common to both vectors. In contrast to E4, there was much less change in PI at E3 (wt PI=1.41 vs. DM PI=1.65) where promoter H3K18/27 acetylation had only a modest effect on transcription (19) (Figure 1b). Similar to E3, the low level of GRO-seq counts at the E2early promoter region showed little difference in PI between wt E1A and DM-E1A (wt PI=0.164 vs. DM PI=0.123) (Figure 1b), suggesting that H3K18/27 acetylation at the E2early promoter primarily promotes Pol2 initiation. Therefore, loss of H3K18/27 promoter acetylation resulted in a smaller defect on promoter-proximal Pol2 pause-release at the E3 and E2early promoters than at the E4 promoter.

### Decreased Pol2 pause-release in the E4 promoter-proximal region correlates with decreased association of SEC subunits CDK9, AF9, and ENL

Phosphorylation of Ser5 on the Pol2 CTD by the CDK7 subunit of TFIIH occurs during transcription initiation, and subsequent CTD-Ser2, NELF, and DSIF phosphorylation by the CDK9 subunit of P-TEFb allows release of Pol2 arrested by NELF binding in the promoter-proximal region, and the transition to productive elongation (6, 7, 21). To characterize these mechanisms on the HAdV-5 genome, we performed ChIP-seq for Pol2 Ser5-P, Pol2 Ser2-P, NELF, and CDK9 in cells expressing wt E1A (Figure 1a). At E2early, Ser 5-P peaked near the TSS and decreased throughout the gene body, a distribution that is typical in yeast which also has short genes with few introns (22), as well as mouse ES cells (21) with the much longer, multi-exon, long intron genes typical of vertebrates. We observed two Ser2-P peaks in the E2early gene body, one just downstream of the TSS, likely indicating paused Pol2. Another Ser2-P peak occurred over the E2early second exon. A small Ser5-P peak was also observed at this position (Figure 2a). These Pol2 peaks may be explained by a reduction in elongation rate over exons, proposed to influence splice site recognition and spliceosome assembly (23, 24). Such a decrease in Pol2 elongation rate over the short E2 second exon would cause an increase in the steady-state level of Pol2 over the exon, potentially leading to the increase in the Pol2 ChIP-seq signal observed over the E2 second exon.

Both CDK9 and NELF peaks occurred at the expected E2early, E3, and E4 pause sites ∼40 bp downstream of the TSSs (Figure 2a). Broad enrichment of Ser2-P and Ser5-P Pol2 also was observed downstream of the E3 and E4 poly(A) sites. Increased Pol2 Ser2-P and Ser5-P downstream from cellular poly(A) sites is observed at most cellular genes in mammalian cells, and is thought to result from a decrease in Pol2 elongation rate following nascent RNA cleavage at the poly(A) site (21).

We next asked if defective paused Pol2 release after activation by DM-E1A was due to decreased recruitment of P-TEFb containing complexes. A large percentage of P-TEFb exists in complex with the 7SK snRNP where its CDK9 kinase activity is inhibited and it is sequestered from chromatin (25–27). Eviction of P-TEFb from the 7SK snRNP enables its integration into complexes with activated CDK9 kinase activity, including the super elongation complex (SEC) and a complex comprised of P-TEFb and BRD4 (28, 29). Integration into these complexes allows active CDK9 to be targeted to promoters and enhancers where it phosphorylates its targets and stimulates paused Pol2 release (10). To determine the effects of H3K18/27ac on SEC and P-TEFb-BRD4 recruitment to early adenovirus genes, we performed ChIP-seq for CDK9, AF9, ENL, and BRD4 on the HAdV-5 genome in infected cells (Figure 2b). Reduced H3K18/27ac in DM-E1A vector-infected cells compared to wt E1A-expressing cells correlated with decreased CDK9, AF9, and ENL association with the early viral promoters and gene bodies compared to cells expressing wt E1A (Figure 2b). Importantly, we did not observe decreases in average AF9 and ENL association with TSSs of most human genes in the same infected cells expressing DM-E1A, demonstrating the specificity of this effect on SEC subunit association at the early viral promoters (Figure 2c). BRD4 association at the E2early, E3, and E4 TSSs changed very little in cells infected with the DM-E1A vector compared to the wt E1A vector, although it was reduced to about 50% the level with wt E1A within the transcription units (Figure 2b). These data suggest that H3K18/27ac facilitates paused Pol2 release at E4 by recruitment of the SEC. In contrast, transcription initiation at E3 requires much less SEC recruitment to achieve a transcription rate near that in control DMSO-treated cells (Figures 1 and 2b).

### CBP/p300 acetyl-transferase activity is required for efficient Pol2 pause-release and recruitment of AF9, ENL, and BRD4 at E4

A-485 is a potent and specific small molecule inhibitor of p300/CBP acetyl transferase activity that competes with acetyl-CoA for binding to the acetyl transferase domain active site (20). Decreased total cell H3K18ac after A-485 treatment in HBTECs was confirmed by western blot (Figure 3a). ChIP-seq for H3K9ac, H3K18ac, and H3K27ac on the HAdV-5 genome in wt E1A vector-infected cells treated with A-485 for 2 h demonstrated inhibition of H3K18/27ac, as expected (20, 30), and slight inhibition of H3K9ac at early viral promoters (Figure 3b). As a measure of the transcription rate of the early viral genes, we assayed pre-mRNA levels using qRT-PCR of intronic RNA in RNA isolated from HBTECs expressing wt E1A treated with 10 µM A-485 or control DMSO vehicle alone for 2h. We observed decreases in E2early and E4 pre-mRNA after A-485 treatment, while E3 pre-mRNA was decreased only moderately (Figure 3c). This result confirms again that E3 transcription is less dependent on promoter region H3K18/27 acetylation than transcription from the E2early and E4 promoters.

**Figure 3:**
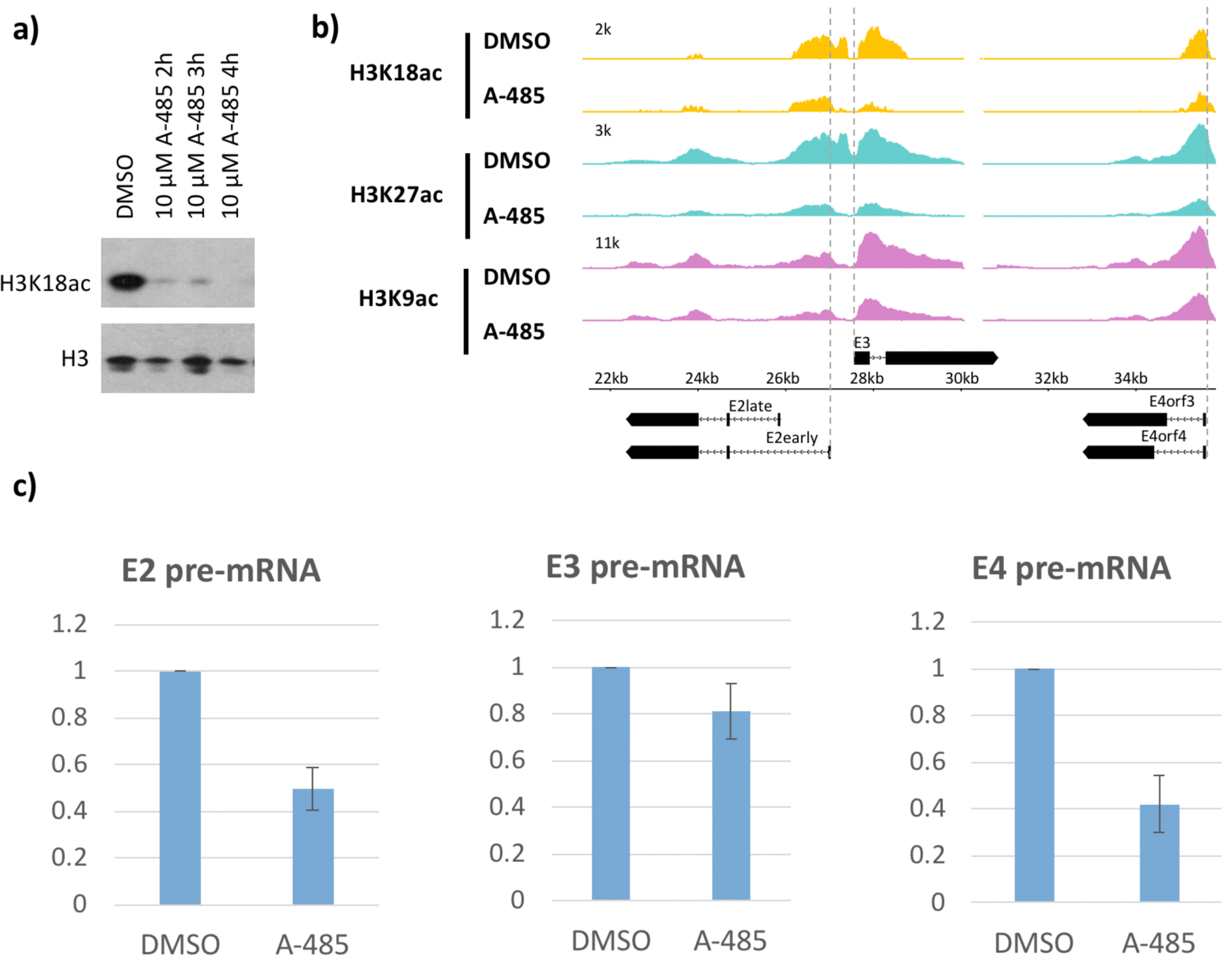
CBP/p300 HAT inhibitor A-485 causes H3 hypoacetylation and decreased early viral gene expression. (a) Western blot for H3K18ac and total H3 in HBTECs treated with 10uM A-485 after 2, 3, or 4 hours. (b) H3K18ac, H3K27ac, and H3K9ac ChIP-seq at early viral promoters in cells treated with 10uM A-485 for 2 hours. (c) qRT-PCR for E2early, E3, and E4 pre-mRNA transcripts in cells treated with 10uM A-485 for 2 hours.

We calculated the pausing indices for transcription of the early viral genes (Figures 4a,b). There was little difference in PI with A-485 treatment at the E2early (DMSO PI=0.13, A-485 PI=0.12) or E3, (DMSO PI=1.20, A-485 PI=1.19) promoters, but a clear increase in PI was observed for E4 (DMSO PI=0.79, A-485 PI=1.13) (Figure 4b). These results indicate decreased release of paused Pol2 after A-485-induced E4 promoter H3K18/27 hypoacetylation. Overall, our results indicate that CBP/p300 HAT activity is necessary for efficient promoter-proximal paused Pol2 release in E4 and are consistent with our results for E4 activation by DM-E1A, where decreased promoter H3K18/27 acetylation also correlated with decreased release of promoter-proximal Pol2 (Figure 1a, bottom).

**Figure 4:**
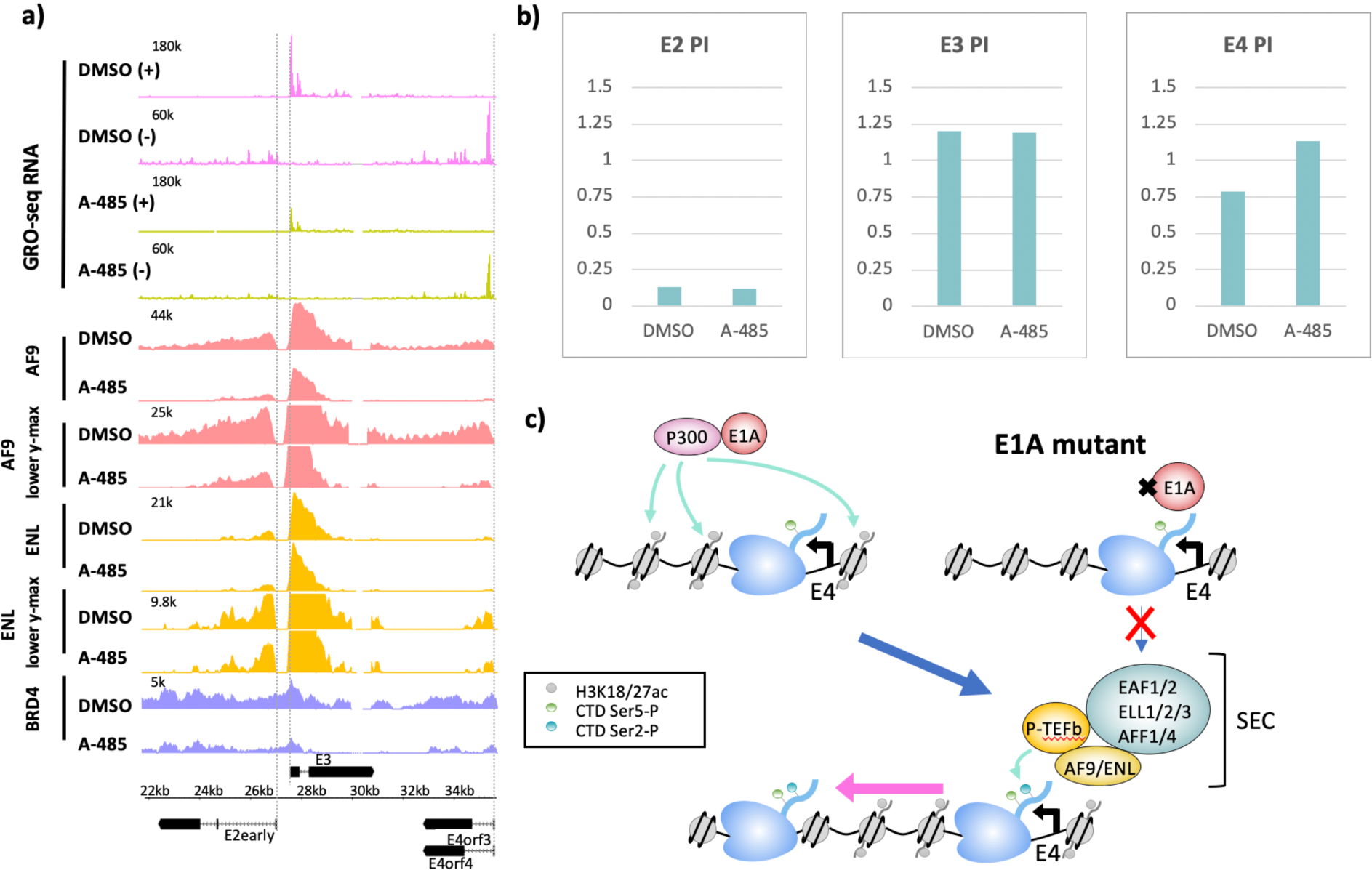
CBP/p300 HAT inhibition by A-485 results in defective Pol2 pause-release and decreased SEC and BRD4 binding at E4. (a) GRO-seq in cells expressing wt E1A treated with DMSO or 10µM A-485 for 2 hours. GRO-seq tracks are plotted with ChIP-seq for AF9, ENL, and BRD4 in cells treated with DMSO or 10 µM A-485 for 2h. Both AF9 and ENL ChIP-seq tracks are shown with two different y-axes. (b) Pause indexes for E2early, E3, and E4 in cells treated with DMSO vs. A-485. (c) Model for regulation of E4 elongation by SEC recognition of CBP/p300-E1A mediated H3K18/27ac.

To determine if inhibition of p300 HAT activity resulted in defective SEC recruitment at E4, we performed AF9, ENL, and BRD4 ChIP-seq in cells infected with the wt E1A vector after DMSO or A-485 treatment. Similar to cells expressing DM E1A, we observed decreases in AF9 and ENL association at the E2early, E3, and E4 promoter regions in cells treated with A-485 (Figure 4a). We also observed decreased BRD4 throughout the transcribed early regions. These data indicate that H3K18/K27 acetylation by CBP/p300 promotes BRD4 and SEC complex association with viral chromatin. This association of BRD4 and SEC complexes with E4 chromatin requires H3K18/27 acetylation by the CBP/p300 acetyl-transferase catalytic domain targeted to the early viral promoters by the interaction between CBP/p300 and the E1A-AD acidic regions (Figure 4c).

### CBP/p300 HAT inhibition by A-485 affects H3 acetylation of cellular chromatin differently at promoters and enhancers

To determine if the effects of H3K18/27ac on Pol2 pause-release at E4 is a general mechanism that also applies to transcription of cellular genes, we shifted our study to the human genome. First, we characterized the changes in H3K18/27ac in HBTECs after 2h treatment with 10 µM A-485. Western blotting demonstrated an extensive decrease in total cellular H3K18ac which approached steady-state by 2 h after addition of A-485, as expected (20, 30) (Figure 3a). Localization of the remaining H3K18ac and H3K27ac was determined by separate ChIP-seq analyses with antibody specific for either H3K18ac or H3K27ac. These results showed that some sites of H3K18/27ac were far more resistant to A-485 treatment than others. Comparing the average signals for H3K18ac and H3K27ac at all TSSs and enhancer peaks (peaks >2.5kb from the nearest TSS), we observed the expected decreases in H3K18ac and H3K27ac by A-485 at enhancer peaks (Figure 5a). However, we observed a surprising <u>increase</u> in the average level of H3K18ac and H3K27ac at all TSSs in cells treated with A-485 (Figure 5a). These observations indicate that homeostatic mechanisms function to maintain H3K18/27ac at promoters when CBP/p300, the principle cellular acetyl transferases for these sites (30–32), are extensively inhibited.

**Figure 5:**
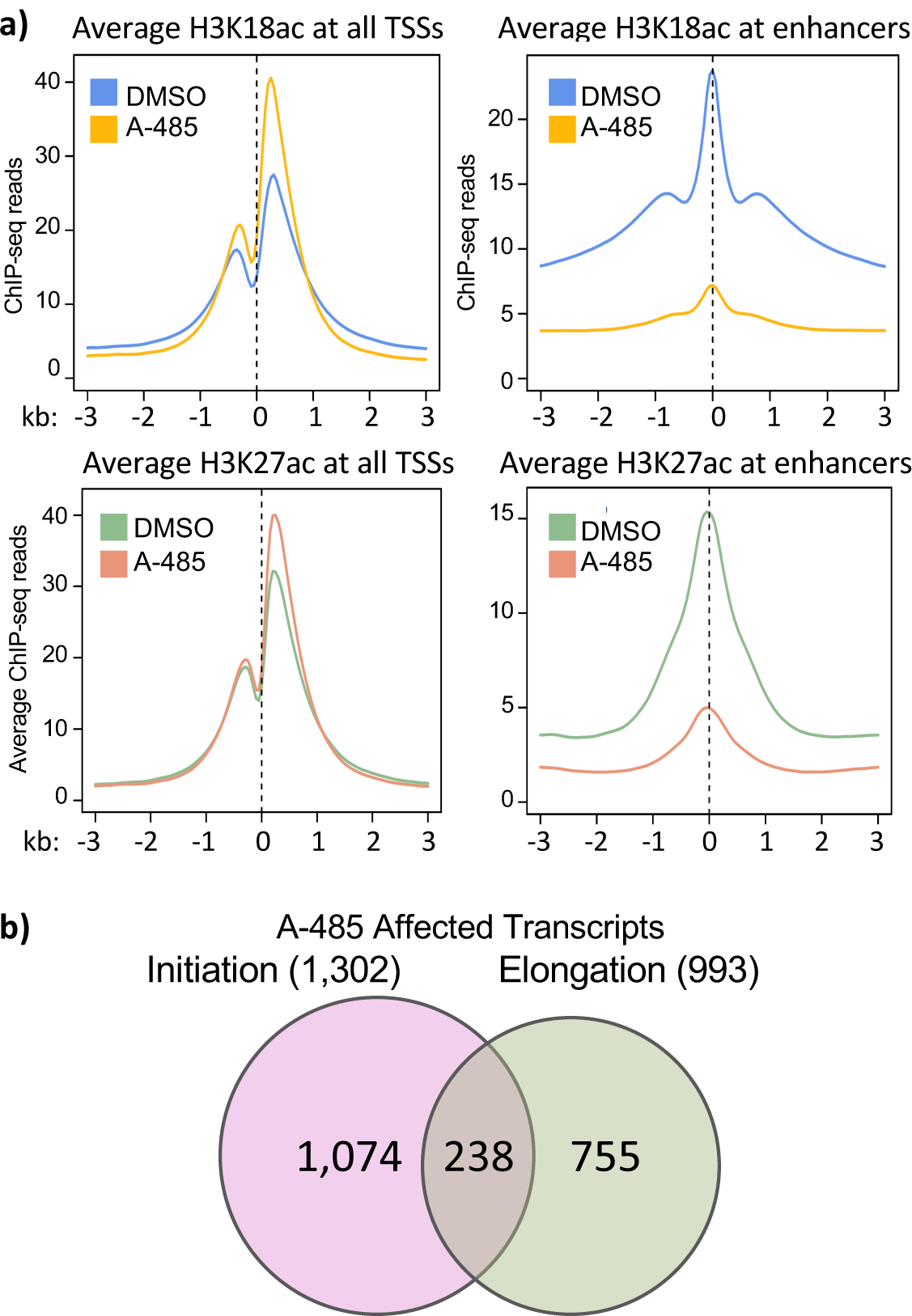
Treatment with A-485 causes different effects on H3K18/27ac at promoters and enhancers and results in defects in both initiation and elongation. (a) Plots of average H3K18ac and H3K27ac signals at all human TSSs and at enhancers in cells treated with A-485 or DMSO. (b) Number of protein coding transcripts with defects in transcription initiation (>2-fold decrease in GRO-seq counts in TSS to +200) and elongation (>2-fold increase in PI).

### A-485 affects cellular genes during both transcriptional initiation and elongation

We were also curious about whether A-485 treatment affected transcription of human genes during both initiation and elongation as we had observed for the HAdV-5 genome, and whether or not there are variations in the effect of A-485 on initiation versus elongation at different human promoters, as observed on the HAdV-5 genome.

GRO-seq reads from control DMSO and A-485 treated cells were aligned to the human genome to determine the fraction of genes affected by A-485 at different stages in transcription. We limited our analysis to protein coding transcription units with active promoters containing at least 20 GRO-seq counts in the promoter region (TSS to +200) and a significant H3K9ac TSS peak (q-value <0.05). Out of 15,768 such active protein coding transcription units, we found 1,302 where initiation was inhibited after 2h A-485 treatment (<50% the GRO-seq counts in the promoter region compared to control DMSO-treated cells), and 993 (6.3%) with defective pause-release after A-485 treatment (>2-fold increase in PI) (Figure 5b). 238 assessed transcription units passed the criteria for both groups, indicating that both transcription initiation and promoter-proximal pause release were reduced by A-485 treatment (Figure 5b).

### A-485 sensitive Pol2 pause-release and SEC recruitment at genes with A-485-induced hypoacetylation at TSSs

First, we consider the 993 protein coding transcription units in which promoter-proximal pause-release was inhibited by A-485 treatment and the resulting loss of H3K18/27ac in their promoter regions. Protein coding transcription units where A-485 treatment caused >2-fold increase in PI are referred to as “2XPI genes.” We plotted the average H3K18, K27, and K9 acetylation ChIP-seq counts near the TSS for all genes and for 2XPI genes (Figure 6a). H3K18ac and K27ac at TSSs for 2XPI genes decreased in response to A-485, as expected for a specific competitive inhibitor of the CBP/p300 acetyl-transferase. But this was in contrast to the surprising *increase* in H3K18 and K27 acetylation on average at the TSSs for all genes in response to A-485. Thus H3K18/27 acetylation in the promoter regions of 2XPI genes was particularly sensitive to CBP/p300 inhibition; whereas, the average H3K18/27 acetylation in the promoter regions for all genes was increased by treatment with A-485 (Figure 6a). H3K9ac, did not change at the TSS in A-485-treated cells in the average plot for all genes, and decreased modestly at TSSs of 2XPI genes after A-485 treatment (Figure 6a).

**Figure 6:**
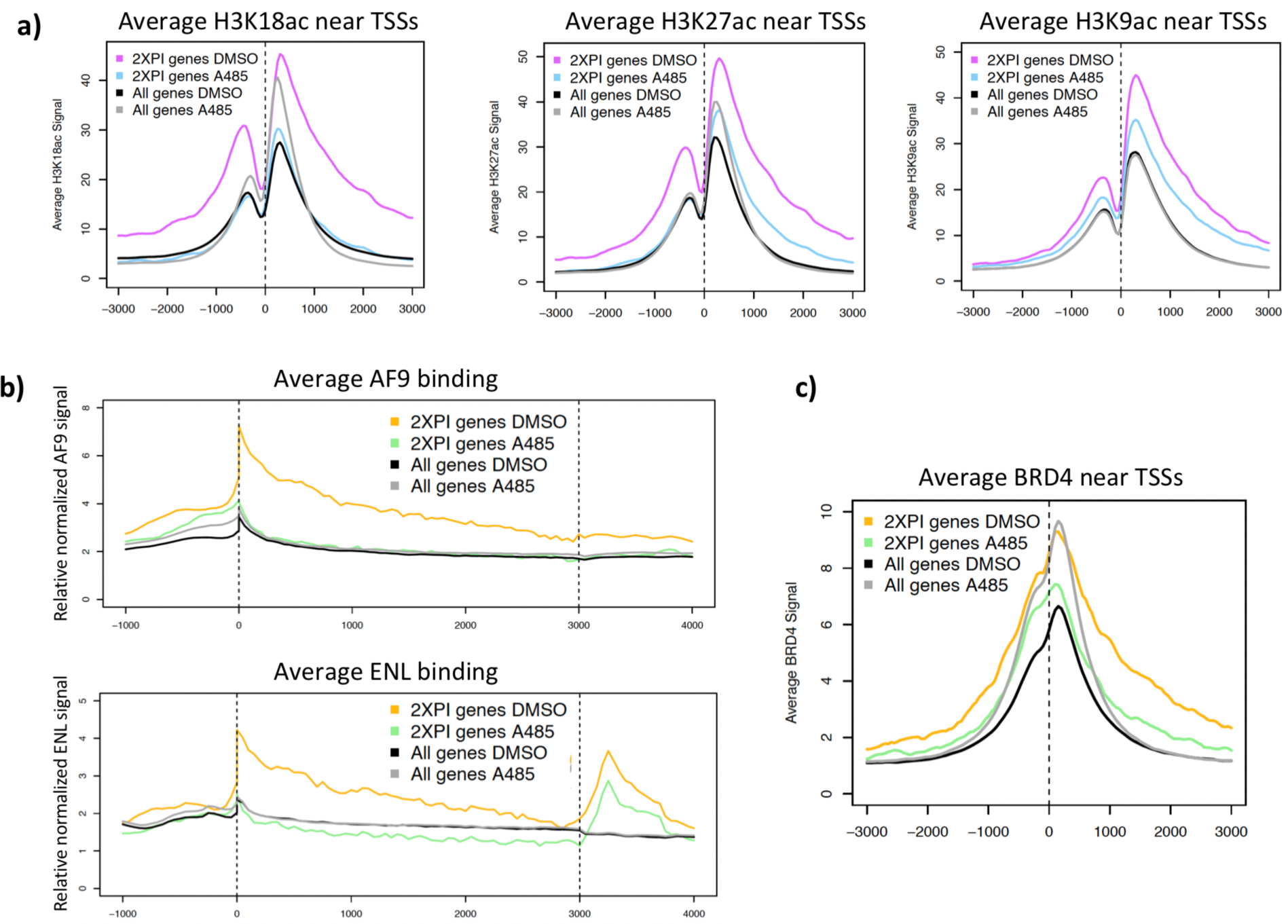
Decreased average H3 acetylation and SEC binding at 2XPI genes. (a) Average H3K18ac, H3K27ac, and H3K9ac near TSSs in all genes and 2XPI genes after DMSO or A-485 treatment. (b) Average AF9 and ENL across all genes and 2XPI genes after DMSO or A-485 treatment. (c) Average BRD4 near TSSs in all genes and 2XPI genes after DMSO or A-485 treatment.

To determine if these decreases in H3 acetylation at 2XPI genes were correlated with decreased SEC component binding, we plotted AF9 and ENL ChIP-seq counts for all genes and for 2XPI genes after A-485 treatment (Figure 6b). Remarkably, AF9 and ENL were highly enriched at TSSs and gene bodies of 2XPI genes. Further, after A-485 treatment, AF9 and ENL association with 2XPI genes fell to the average level for all genes. Thus, genes with an increase in PI after A-485 treatment were very highly enriched for association of SEC complexes throughout their transcription units. BRD4 association near the TSS of 2XPI genes was reduced by A-485 (Figure 6c) but to a far less extent than AF9 and ENL reduction (Figure 6b).

### H3K9ac is sufficient for BRD4 enhancer binding which stimulates pause release at nearby genes

It was evident that BRD4 peaks were enriched at enhancers. 56% of identified BRD4 peaks (14,877 out of 26,534 (see Methods)) were >2.5kb from the nearest TSS in control cells treated with DMSO. Of these distal peaks, 84% overlapped with peaks of H3K27ac, indicating that these peaks were primarily at enhancers. We subsequently clustered all enhancers associated with BRD4 based on whether or not BRD4 association decreased to <50% of DMSO control after A-485 treatment for 2h. There were 7,554 peaks where BRD4 decreased to <50% of control after A-485 treatment (referred to as “A-485 sensitive” enhancers; Figure 7a bottom) and 6,324 peaks where BRD4 association remained unaltered or was not reduced to <50% of control (“A-485 resistant”; Figure 7a, top). As shown in Figure 7a, decreased BRD4 binding after A-485 treatment correlated with decreased H3K9ac. When BRD4 binding was unaffected or modestly affected by A-485 treatment, the decrease in H3K9ac was minimal. These observations suggest that acetylation at H3K9 is sufficient for BRD4 association at enhancers.

**Figure 7:**
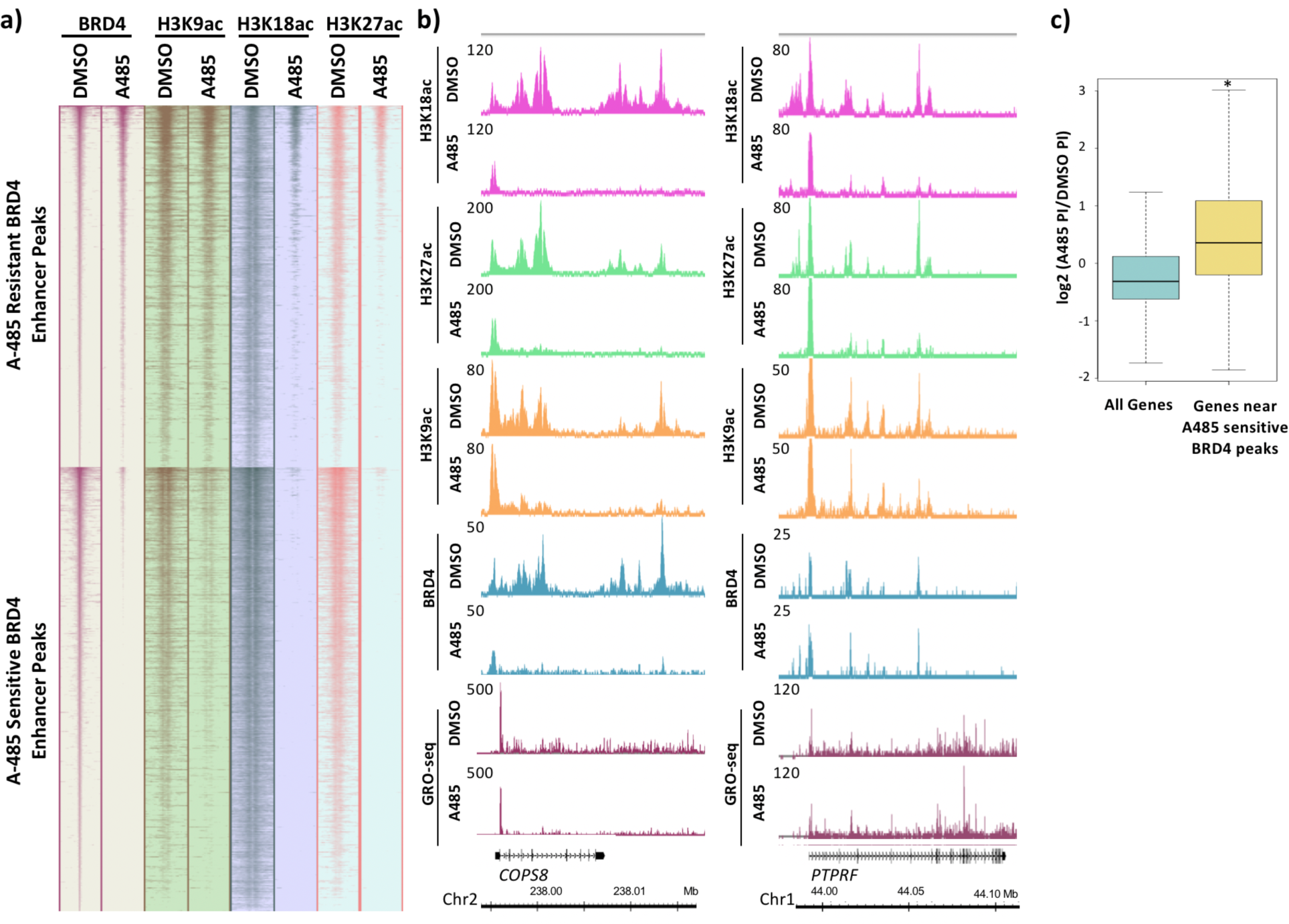
BRD4 enhancer binding stimulates pause-release at nearby genes. (a) Heatmaps of BRD4, H3K9ac, H3K18ac, and H3K27ac ChIP-seq data. BRD4 enhancer peaks are divided into those which are A-485 resistant (top cluster) or A-485 sensitive (bottom cluster). (b) Gene browser plots of ChIP-seq data for the indicated histone modifications and BRD4, and GRO-seq counts for regions including the *COPS8* (left), and *PTPRF* (right) genes. (c) Boxplots comparing the change in PI after A-485 treatment (log2 (A-485 PI/DMSO PI)) for all genes vs. genes near A-485 sensitive enhancer BRD4 peaks.

We next asked if BRD4 enhancer association correlated with the extent of Pol2 pause-release in the promoter proximal region of nearby genes. For example, *COPS8* is a gene with downstream proximal enhancers that have A-485-sensitive BRD4 association (Figure 7b). At these enhancers there was decreased H3K18/27ac, H3K9ac, and BRD4 in A485-treated cells (Figure 7b). This correlated with only a modest decrease in the GRO-seq reads at the Pol2 pause site (∼30%, Figure 7b), and therefore, an ∼30% decrease in the amount of Pol2 that had initiated transcription at the *COPS8* TSS in this population of cells, compared to control DMSO-treated cells. But A-485 caused a larger decrease in GRO-seq reads downstream from the *COPS8* promoter-proximal pause site (Figures 7b,S2). *NDRG1* is an example of another gene proximal to enhancers with A-485-sensitive BRD4 association. Similar to *COPS8*, A-485 treatment caused a decrease in release of promoter-proximal paused Pol2, but little decrease in Pol2 initiation near the pause site (Figure S3). These results indicate that A-485 inhibits *COPS8* and *NDRG1* transcription primarily during the release of Pol2 from the major promoter-proximal pause site.

In contrast to *COPS8* and *NDRG1, PTPRF* is a gene with A-485-resistant BRD4 association at nearby enhancer regions in its introns (Figure 7b). A-485 treatment reduced H3K18/27ac but not H3K9ac or BRD4 association at these enhancers. This correlated with only a modest decrease in the GRO-seq reads at the Pol2 pause site (to ∼70% the level in control DMSO-treated cells, Figure 7b), and therefore, an ∼30% decrease in the amount of Pol2 that had initiated transcription at the *COPS8* TSS in this population of cells, compared to control DMSO-treated cells. These results indicate that *PTPRF* is not regulated by H3K18/27ac during elongation, and instead suggest that H3K18/27ac primarily regulates transcription initiation of the *PTPRF* gene. Similarly, *PAG1*, encoding a transmembrane adaptor protein that organizes membrane-proximal signaling complexes in lipid rafts, is another example of a gene that showed A-485 inhibition of Pol2 pause release in the promoter proximal region (Figure S4). Thus, *PTPRF* and *PAG1* appear to be regulated by promoter-proximal H3K18/27ac stimulation of promoter-proximal pause release, similarly to the HAdV-5 E4 promoter (Figure 1a, bottom). Alternatively, *CSF3* is an example of a gene where promoter region H3K18/27 hypoacetylation greatly inhibited Pol2 initiation (Figure S5), as for the HAdV-5 E2early promoter. *TRIB1* (Figure S6) is an example of a gene where A-485 and the resulting promoter region H3K18/27 hypoacetylation inhibited both initiation (to ∼50% the level in control DMSO-treated cells) and elongation passed the pause site.

Comparing the *COPS8* and *PTPRF* genes, the major difference in H3 acetylation in response to A-485 was at H3K9 in their associated enhancers (Figure 7b). H3K9ac of the *PTPRF* intronic enhancer regions was only minimally reduced by A-485 treatment and correlated with A-485-resistant BRD4 association with the intronic enhancer (Figure 7b). Whereas at the *COPS8* and *NDRG1* genes, A-485 treatment inhibited H3K9ac at the downstream enhancers, and this loss of enhancer H3K9ac correlated with reduced BRD4 association with these enhancers (Figure 7b, S3). Reduced BRD4 association with these intronic enhancers of *COPS8* and *NDRG1* correlated with decreased pause release from their promoter-proximal pause sites after treatment with A-485 (Figure 7b). This correlation between A-485-sensitive BRD4-enhancer association and efficient Pol2 pause-release in the promoter-proximal region was observed broadly. When we compared the difference in PIs after A-485 treatment, we observed a significant increase in PI distribution for genes near A-485-sensitive enhancer BRD4 peaks compared to all genes (Figure 7c).

## Discussion

There is a substantial base of knowledge establishing a correlation between histone N-terminal tail lysine acetylation and transcriptional activity. However, understanding of the mechanisms underlying this correlation remains incomplete. In addition to regulating PIC assembly and Pol2 initiation, our results support a mechanism by which histone H3 N-terminal tail acetylation in the promoter-proximal region regulates Pol2 release from promoter-proximal pause-sites in a subset of adenovirus and primary human airway epithelial cell promoters.

### H3K18/27ac by CBP/p300 stimulates BRD4 and SEC-association and promoter-proximal Pol2 pause-release

Initially we observed that super-elongation complex (SEC) recruitment through association with promoter region histone H3 acetylated at K18 and K27 stimulates paused Pol2 release at the HAdV-5 E4 promoter (Figure 1a). To do this study, we used a multi-site E1A mutant (DM-E1A) defective for binding CBP/p300 by the E1A activation domain (19), and defective for stimulating promoter H3K18/27ac at viral early promoters (19). GRO-seq studies following infection of primary airway epithelial cells with Ad5 vectors expressing wt or DM-E1A revealed that the E4 pausing index increased when the DM-E1A failed to stimulate H3K18/27ac at the E4 promoter. This result suggests that promoter H3K18/27ac contributes to paused Pol2 release at the E4 promoter.

We observed a correlation between defective paused Pol2-release at the viral E4 promoter and decreased association of SEC subunits CDK9, AF9, and ENL, indicating that H3K18/27ac is necessary for maximal SEC recruitment and Pol2 pause-release in the E4 promoter region. This is similar to a proposed function of H3K9ac as a binding site for AF9 and ENL, thereby promoting paused Pol2 release by directly recruiting the SEC (15). SEC recruitment at E4 by H3K18/27ac may be due to interactions with acetyl-lysine-binding YEATS domains present in the AF9 and ENL SEC subunits (14, 15), but other SEC components may also contribute. For example, the SEC was reported to be recruited to chromatin with H3K27ac through an interaction with the C-terminus of the central scaffold protein AFF4 at the TSS of the estrogen receptor 1 gene (*ESR1*) in cultured breast cancer cells (33).

Defective pause-release and decreased SEC recruitment at E4 also were observed in wt E1A-expressing cells when CBP/p300 acetyl-transferase activity was inhibited by the competitive inhibitor A-485 (19) (Figure 4A). These results indicate that CBP/p300 HAT activity is necessary for maximal promoter-proximal paused Pol2 release at E4. However, these data do not rule out the possibility that other factors that associate with H3K18/27ac also contribute to Pol2 pause-release at E4. For example, BRD proteins have several BRD domains, some of which bind acetylated lysines with moderate affinity, potentially participating in cooperative protein binding to a region of chromatin with multiple acetylated lysines (29, 34, 35). Additionally, the Mediator complex subunit MED26 is known to recruit the SEC after dissociation of the mediator from TFIID (36, 37). Therefore, it is also possible that additional consequences of promoter proximal H3K18/27 hypoacetylation, such as reduced association with MED26-containing mediator complexes, also contribute to decreased Pol2 promoter-proximal pause release at E4 after activation by DM-E1A.

### A consensus TATA-box overcomes transcription inhibition by promoter H3K18/27 hypoacetylation

It is interesting to note that the sensitivity of HAdV-5 early region transcription to promoter H3K18/27 hypoacetylation correlated with the similarity between their TATA-box sequences and the consensus TATA-box sequence. This likely results in higher affinity of TBP for the TATA-boxes of the genes resistant to H3K18/27 hypoacetylation, than for the sensitive genes. E3 transcription is the most resistant of the HAdV-5 early regions to promoter hypoacetylation (19). The E3 TATA-box is a good match to the consensus TATA-box sequence TATA[A/T]A[A/T][A/G]. It’s sequence, cg<u>**TATAA**C**T**C</u>ac (central eight base pairs contacted directly by human TBP (38) shown capitalized, and matches to the TATA-box consensus sequence shown bold). The match to the consensus TATA-box is particularly good in the 5’-half of the TATA-box which makes more contacts with TBP than the 3’-half of the TATA-box and is more highly conserved than the 3’-half (38). In contrast, E2early is the most sensitive early region to promoter H3K18/27 hypoacetylation (19), and it’s TATA-box (cc**T**TA**A**G**A**GTca) has the lowest match to the consensus TATA-box. This probably results in a lower affinity of TBP for the E2early compared to the E3 TATA-box. Promoter H3K18/27 hypoacetylation at the E2early promoter in response to A-485 treatment greatly inhibited transcription initiation as indicated by the decrease in GRO-seq counts at the E2early TSS and gene body (Figure 4a).

The HAdV-5 E4 promoter has a symmetrical TATA-box (cc**TATATATA**ct) (39) that is a perfect match to the consensus TATA-box in the eight base pairs that interact directly with human TBP (38). Again, GRO-seq showed that despite E4 promoter H3K18/27 hypoacetylation (Figure 1a), there was little if any defect in Pol2 initiation and elongation to the promoter-proximal pause site (Figures 1a highlighted and 2a). Thus, compared to the E2early promoter with a non-consensus TATA-box, promoter H3K18/27 hypoacetylation had a much smaller effect on transcription initiation at the E3 and E4 promoters with consensus TATA-boxes that are probably bound by TBP with higher affinity than the E2early TATA-box.

The principle effect of E4 promoter H3K18/27 hypoacetylation was on promoter-proximal pause release, causing a decrease in transcribing Pol2 downstream from the promoter revealed by low GRO-seq counts in the gene body (Figure 1a). This correlated with lower BRD4, CDK9 and SEC subunit association throughout the gene body after activation by DM-E1A compared to wt E1A (Figure 2b).

### A subset of cellular promoters requires H3K18/27 acetylation by CBP/p300 for maximal promoter-proximal Pol2-release

Analysis of the ChIP-seq and GRO-seq data for cellular chromatin from cells treated with A-485 established that regulation of Pol2 pause-release and SEC recruitment by promoter region H3K18/27ac also occurs at a small fraction of cellular promoters. With A-485 treatment, we observed the expected decreases in H3K18/27ac at enhancers, along with an intriguing <u>increase</u> in the average H3K18/27ac at TSSs of all genes. This indicates that the dynamics of HAT and/or HDAC activities in response to A-485 differs at promoters versus enhancers. A-485 also resulted in defects in pause-release at ∼6% of active promoters in primary respiratory epithelial cells. These promoters are similar in that promoter region acetylation was decreased after A-485 treatment, as opposed to the increase in average promoter H3K18/27ac at all genes (Figure 6a).

### Higher rate of H3K18/27 acetylation at promoters compared to enhancers

The steady-state level of H3K18/27ac on any specific nucleosome is determined by the relative rates of its acetylation and de-acetylation (30, 40). A-485 inhibits CBP/p300 acetyl transferase activity by competing with acetyl-CoA for binding to the enzyme’s active site (20). No evidence for inhibition of a histone deacetylase by A-485 was detected (20), and is very unlikely given the highly specific interactions of A-485 with the CBP acetyl-transferase domain (20). Consequently, since it seems unlikely that A-485 directly increases the rate of H3 deacetylation, the decrease in average enhancer H3K18/27ac in A-485 treated cells to one-third the level in control DMSO-treated cells (Figure 5a) suggests that the rate of H3K18/27 acetylation at enhancers in cells treated with A-485 for 2h or more (Figure 3a) was reduced to one-third of the normal rate in control DMSO-treated cells.

In striking contrast to this expected decrease in H3K18/27ac at enhancers, at promoters the average H3K18/27ac <u>increased</u> during treatment with this specific inhibitor of the CBP/p300 acetyl-transferase activity. This result indicates that the rate of H3K18/27 acetylation at promoters is ∼4 to 5-fold higher than at enhancers. These results also suggest that there is an uncharacterized homeostatic mechanism that maintains promoter region H3K18/27ac in the face of extensive inhibition of the known lysine acetyl transferases that acetylate these sites, the closely related CBP and p300 (18). It is possible that the difference in the effects of A-485 on the rates of promoter versus enhancer acetylation by CBP/p300 is due to differences in nucleosome density or the density of other proteins at promoters versus enhancers that restrict the diffusion of the 536 Da drug molecule to the CBP/p300 active site. However, it seems unlikely that diffusion of A-485 molecules would be greatly restricted by nucleosomes that are ∼400 times larger than the drug and irregularly packed into disordered chains of “beads on a string” nucleosomes with different particle and linker DNA arrangements in interphase nuclei (41). Consequently, the resistance of H3K18/27ac at TSSs to A-485 in living cells probably results from an ∼four to five-fold faster rate of H3K18/27 acetylation by CBP/p300 at promoters than at enhancers and most other locations in the genome, on average. This is the result expected if transient interactions between the activation domains of activators bound to their cognate DNA-binding sites in enhancers increase the local concentration of CBP/p300 in promoter regions.

BRD4 contains two bromodomains that bind acetylated lysines (42, 43). The C-terminal portion of BRD4 binds P-TEFb and is thought to recruit it to hyperacetylated genomic regions to stimulate elongation (22). BRD4 has been shown to associate with promoters and enhancers and to act as a histone chaperone to facilitate elongation of both protein coding and enhancer RNAs (10). By clustering BRD4 enhancer peaks into A-485-sensitive and -resistant groups and correlating these data with H3K9ac and H3K18/27ac association, we conclude that H3K9ac is sufficient for BRD4 recruitment at enhancers in the absence of H3K18/27ac. Additionally, GRO-seq in cells treated with A-485 revealed a correlation between decreased BRD4 enhancer association and defective release of paused pol2 from nearby promoters. The mechanism by which this occurs is likely through direct promoter-enhancer interactions facilitated by long-range chromatin interactions (36). Another possibility is that transcription of enhancer RNAs (eRNAs) stimulated by BRD4 stimulates paused Pol2 release by promoting NELF release (44).

Our results suggest a model in which histone H3 acetylation is essential for maximal paused Pol2 release at the HAdV-5 E4 promoter. By analyzing H3 acetylation, SEC subunit chromatin association, and pol2 pausing on the human genome, we establish that this is mechanism that applies to ∼1000 active human promoters in primary airway epithelial cells. Additionally, the identification of BRD4 enhancer peaks that were either sensitive or resistant to A-485 treatment presented an opportunity to study the effects of elongation factor association with enhancers, on elongation. Interestingly, we found that H3K9ac is sufficient for BRD4 binding at enhancers and that BRD4 enhancer binding is correlated with decreased Pol2 pausing and increased productive elongation. Taken together, our results draw interesting causal links between histone H3 acetylation and regulation of Pol2 elongation as well as initiation.

## Materials and Methods

### Ad5 mutant vectors

Ad5 mutant vectors expressing wt E1A and DM E1A were constructed as previously described (19).

### Cell culture

Human bronchial/tracheal epithelial cells (HBTEC; catalog number FC-0035, lot number 02196; Lifeline Cell Technology) were grown at 37°C in a BronchiaLife medium complete kit (LL-0023; Lifeline Cell Technology) in a 5% CO2 incubator until they reached confluence. Cells were then incubated 3 days more without addition of fresh medium and were infected for 12 hours with the indicated HAdV-5 mutants in the conditioned medium. A-485 (MedChemExpress) was added to a final concentration of 10 µM, or the same volume of DMSO vehicle was added, and cells were incubated for an additional 2 h.

### GRO-seq

Cells were harvested and incubated in swelling buffer (10 µM Tris-HCl, 2 mM MgCl_2_, 3 mM CaCl_2_). Nuclei were isolated with lysis buffer (10 µM Tris-HCl, 2 mM MgCl_2_, 3 mM CaCl_2_, 10% glycerol, 1% NP-40). Nuclear run-on was performed at 30°C for 7 min in 10 mM Tris-HCl pH 8, 5 mM MgCl_2_, 300 mM KCl, 1 mM DTT, 500 µM ATP, 500 µM GTP, 500 µM Br-UTP, 2 µM CTP, 200 U/ml Superase In RNase Inhibitor (Invitrogen), and 1% Sarkosyl. Nuclear RNA was isolated with Trizol (Invitrogen). DNAse treatment was performed with Turbo DNA-free kit (Invitrogen). RNA was purified with Micro Bio-Spin P-30 Gel Columns (Bio-Rad), fragmented with RNA Fragmentation Kit (Invitrogen), and treated with 10 units RppH (NEB) and 30 units T4 PNK (NEB). RNA immunoprecipitation was performed with Anti-BrU-conjugated agarose beads (Santa Cruz Biotechnologies). Library preparation was performed with TruSeq Small RNA Library Preparation Kit (Illumina). GRO-seq reads were aligned with HISTAT2 software to Ad5 and human (hg19) genomes and normalized to the number of reads aligned to hg19. Pause indexes (TSS to +200 counts)/(+201 to TTS counts) were calculated using HTSeq software.

### qRT-PCR

Total RNA extracted from HTBECs using a PureLink RNA minikit (Ambion) was reverse transcribed with random hexamer priming using Superscript III (Invitrogen). RNA was treated with DNase I with Turbo DNA-free kit (Ambion). Quantitative reverse transcription-PCRs (qRT-PCRs) were carried out with the Applied Biosystems 7500 real-time PCR system with FastStart universal SYBR green master mix (Roche). All values were normalized to 18S RNA levels.

### ChIP-seq

Preparation of cross-linked HBTEC chromatin, sonication, and immunoprecipitation was as described in reference (32). Sequencing libraries were constructed from 1 ng of immunoprecipitated and input DNA using the KAPA Hyper Prep kit (KAPA Biosystems) and NEXTflex ChIP-seq barcodes (Bio Scientific).

### Data Analysis of ChIP-seq

ChIP-seq libraries were sequenced using HiSeq 4000 or NovaSeq 6000. For analysis on the Ad5 genome, sequence tags were aligned using Bowtie2 software and normalized to the following formula: (number of Ad5-aligned reads in the input sample/number of human-aligned reads in the input sample) × (number of Ad5-aligned reads in the ChIP sample). For analysis on the human genome, reads were mapped to the hg19 human genome reference using Bowtie2 software. Only reads that aligned to a unique position in the genome with no more than two sequence mismatches were retained for further analysis. Duplicate reads that mapped to the same exact location in the genome were counted only once to reduce clonal amplification effects. MACS2 software was used for peak calling (q-value < 0.05 were considered significant). The total counts of the input and ChIP samples were normalized to each other. Samples were normalized for equal number of uniquely mapped reads. The input sample was used to estimate the expected counts in a window. Wiggle files were generated using a custom algorithm and present the data as normalized tag density as seen in all figures with genome browser shots. Metagene plots displaying normalized average relative ChIP-seq signals were generated using CEAS software.

### Antibodies

Antibodies included H3K18ac (814), prepared and validated as described previously (45), H3K9ac (07-352; Millipore), H3K27ac (39133; Active Motif), H3 (ab10799, Abcam), AF9 (GTX102835, Genetex), BRD4 (A301-985A50), NELF TH1L D5G6W (12265S, Cell Signaling), Pol2 Ser2-P 31Z3G (13499, Cell Signaling), Pol2 Ser5-P D9N5I (13523, Cell Signaling), and CDK9 C12F7 (2316, Cell Signaling).

