## Supplemental Figures for "Promoter-specific changes in initiation, elongation and homeostasis of histone H3 acetylation during CBP/p300 Inhibition"

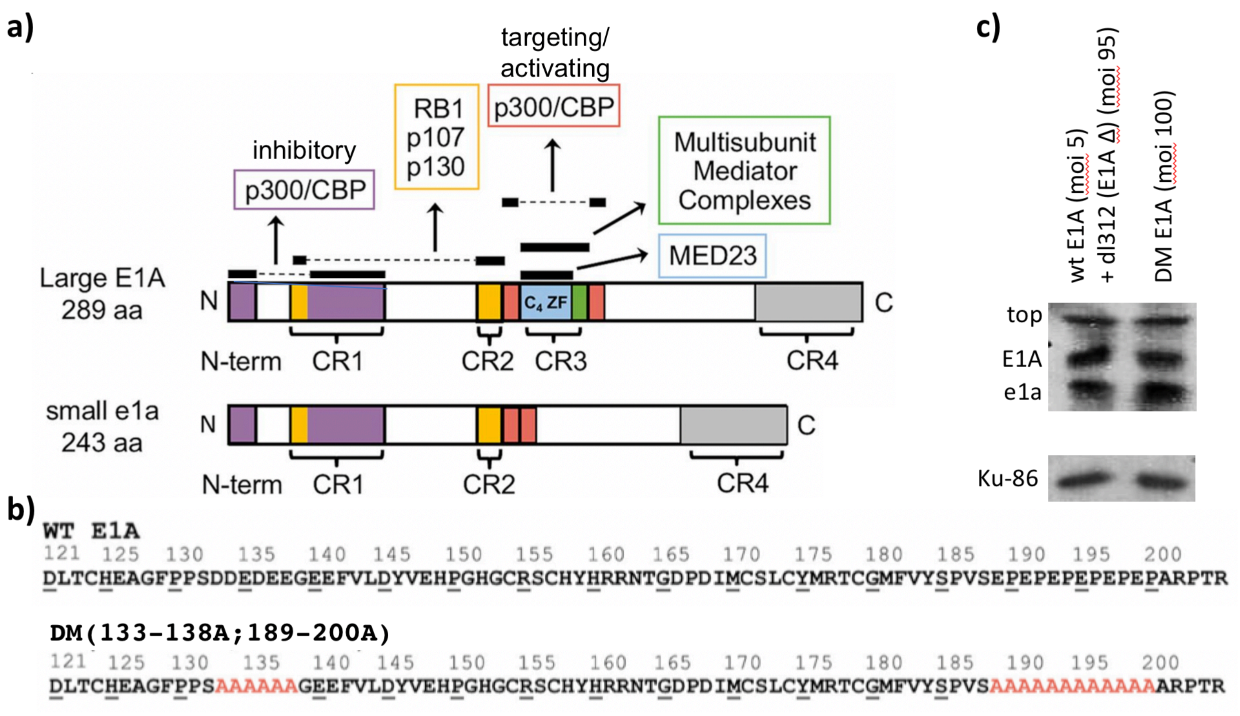


***Figure S1: Map of regions of Large and small E1A Bound by the indicated host cell nuclear proteins:*** **(a)** For both Large and small E1A, the N-terminal ~15 aa residues plus the portion of CR1 diagrammed in purple (aa 54-82 (numbers from Large E1A)) are intrinsically disordered protein (IDP) regions that fold and are bound by the CBP/p300 TAZ2 domain (Ferreon et al. 2009), inhibiting CBP/p300 histone acetyl transferase activity (Borrelli et al (1984), Ferrari et al (2014) and references therein). The N-terminal portion of CR1 (aa 37-49) and CR2 (aa 121-129, both orange) are bound by the “pocket” domains of RB-family proteins (Liu X, Marmorstein R (2007); Ferreon et al. (2009)). p300/CBP are also bound by two acidic regions in Large E1A flanking, CR3 (red), but not the same amino acids when they are juxtaposed in small e1a (Pelka P et al (2009), Hsu E et al (2018)). This interaction between CBP/p300 and the Large E1A activation domain (aa 133-205) activates acetylation of H3K18 and H3K27 at early viral promoters by CBP/p300 (Hsu E (2018). The region diagrammed in blue includes a C4-Zn-finger (Culp et al. (1988) that binds the MED23 mediator subunit (Boyer et al (1999), and the seven amino acid region diagrammed in green in Large E1A that is perfectly conserved in over 40 primate adenoviruses (Avaakumov N et al 2004) and is required for Large E1A binding to multi-subunit mediator complexes (Hsu et al. (2018) and for activation of transcription from the early viral promoters (Boyer et al (1999). CR4 the C-terminal half of the E1A proteins are bound by additional host proteins not discussed here.

**(b)** Sequences of CR3 encoded in the 13S E1A mRNA for the Large E1A protein isoform. Mutant amino acids in DM-E1A are shown in red.

**(c)** Western blot with mAb M58 that binds the same epitope in the N-terminal half of Large and small E1A. This western blot shows that at 12 h post-infection (p.i.), DM-E1A is expressed at the same concentration in cells infected with the DM-E1A vector as wt E1A in cells infected with the wt E1A vector at an m.o.i. = 5 and the E1A deletion mutant *dl312* at an m.o.i. = 95.

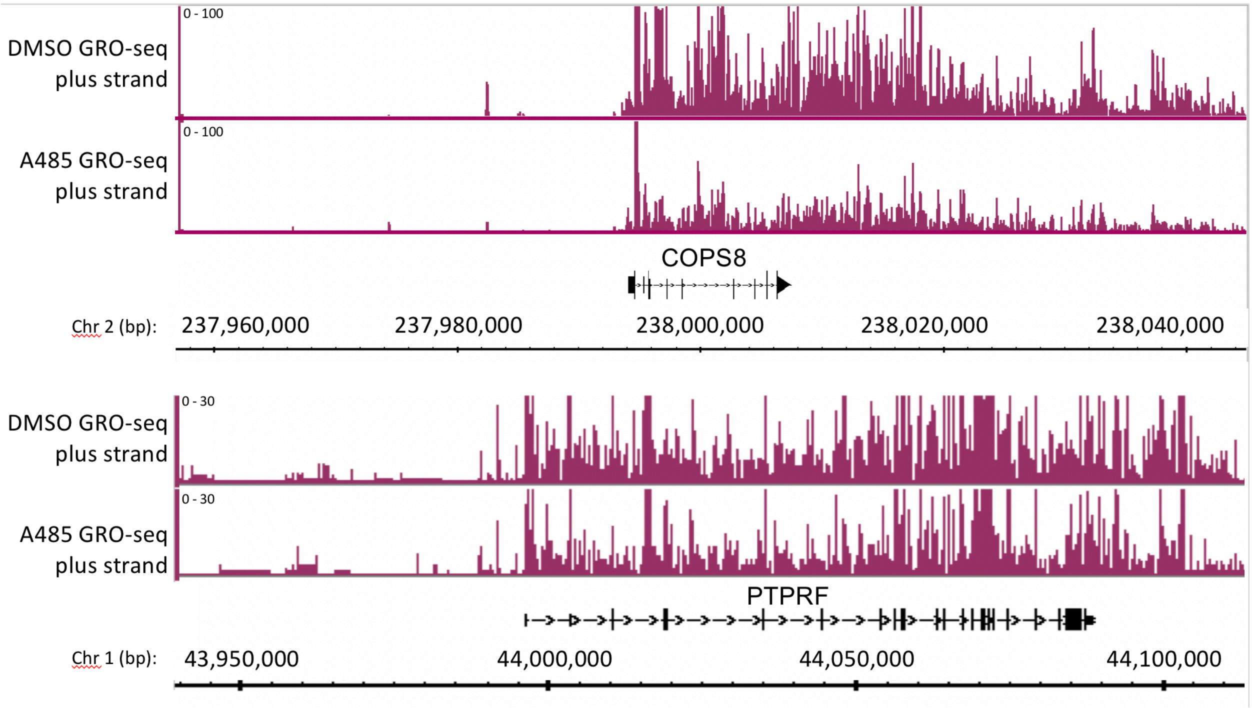


***Figure S2:*** GRO-seq counts plotted with low $y_{max}$ to show the relative level in the gene bodies of *COPS8* and *PTPRF* in response to A-485.

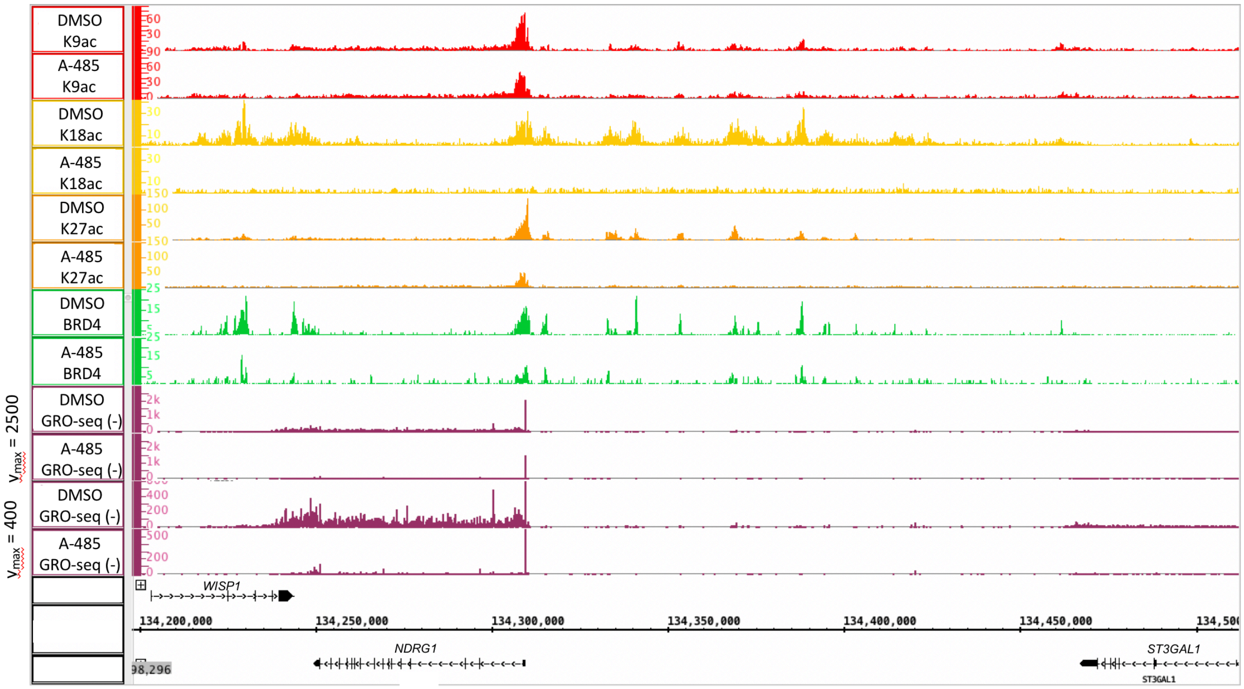


***Figure S3:*** GRO-seq counts plotted with high and low $y_{max}$ to show the relative level of paused Pol2 near the TSS in A-485-treated and control DMSO-treated cells (seen in plot with y max = 2,500), and in the gene body of the *NDRG1* gene (best observed at y max = 400).

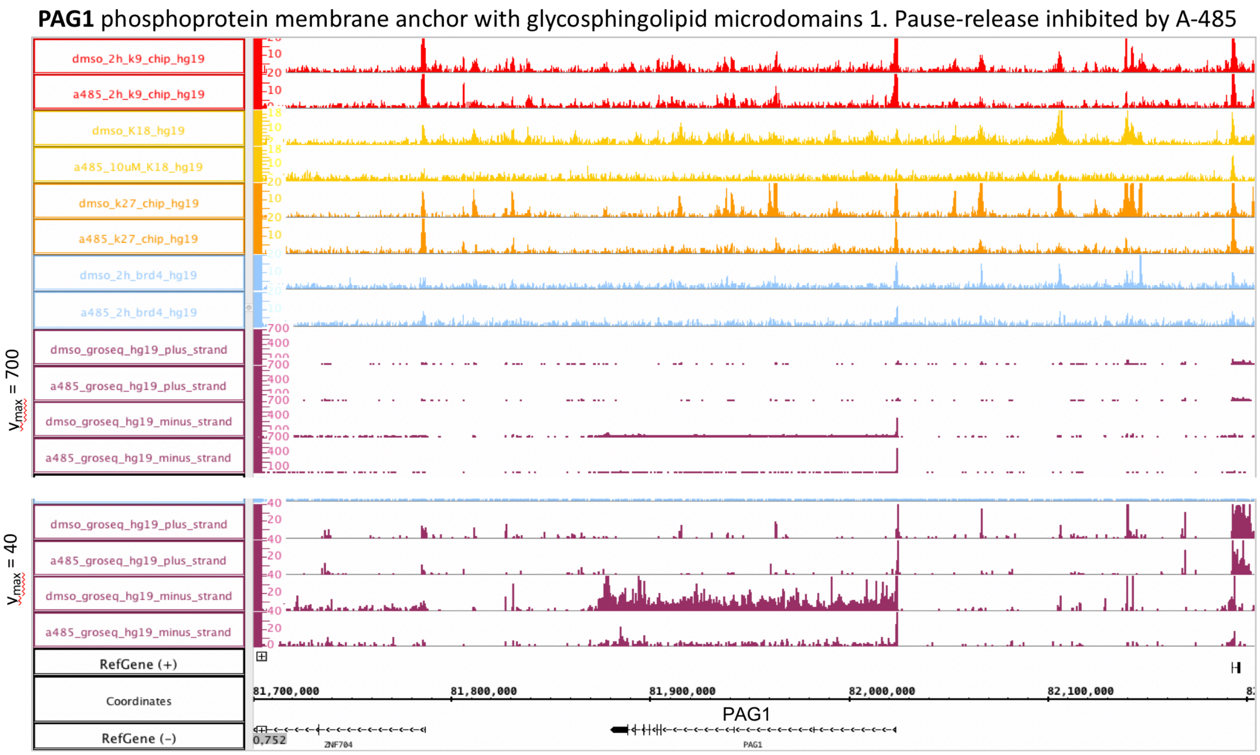


***Figure S4:*** A-485 inhibited Pol2 release from the *PAG1* promoter-proximal pause site, but not Pol2 initiation and transcription to the *PAG1* pause site.

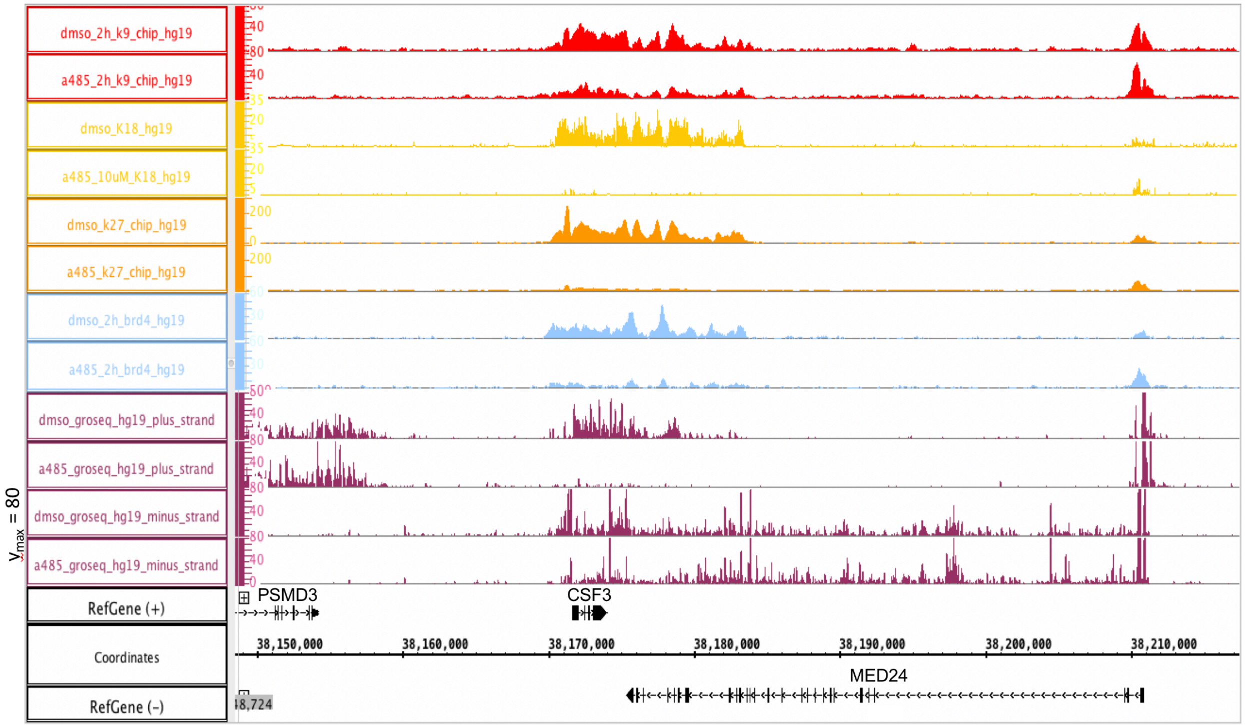


***Figure S5:*** A-485 inhibition of transcription initiation of *CSF3,* but not *PSMD3* to the left also transcribed from the plus strand, or *MED24* to the right, transcribed from the minus strand.

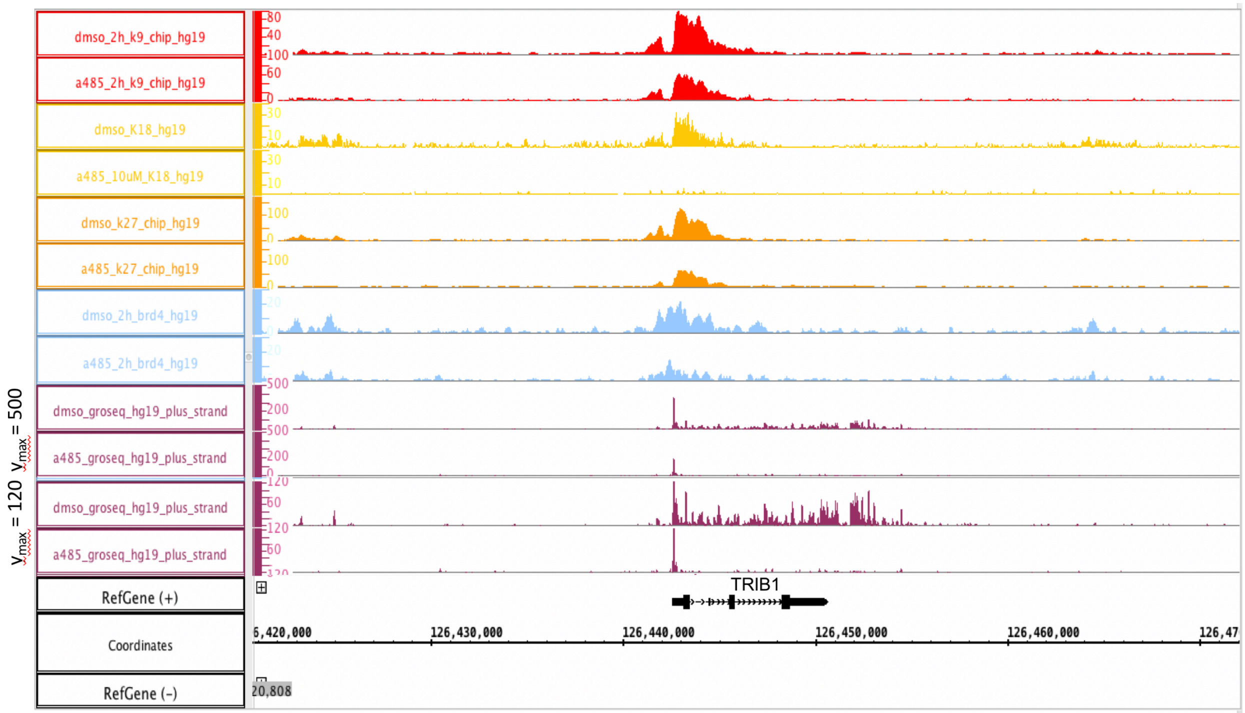


***Figure S6:*** A-485 inhibited Pol2 initiation and transcription to the pause site of *TRIB1* to ~50% the level in DMSO-treated control, and also nearly eliminated Pol2 pause-release and transcription past the pause site.
